## Supplementary data for "A transient protein folding response targets aggregation in the early phase of TDP-43-mediated disease"

**SUPPLEMENTARY MATERIAL**

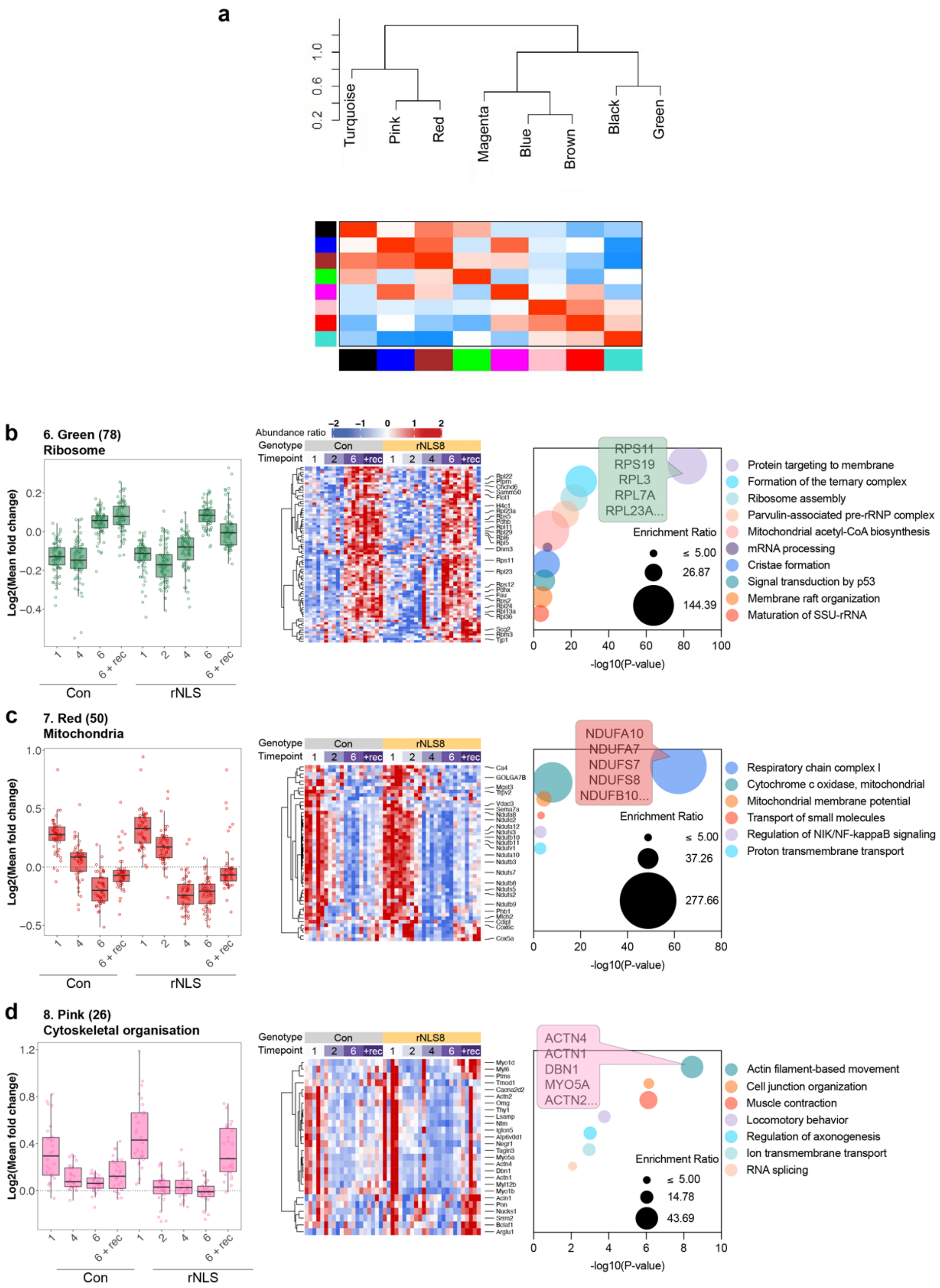


**Supplementary Figure 1. Weighted correlation network analysis reveals 8 defined modules in proteomics of control and rNLS8 mice, of which three are age dependent. a,** Module eigenprotein network showing hierarchical clustering and heatmap of eigenprotein correlation adjacency table. Red in the heatmap represents more closely correlated modules. **b-c,** green, red, and pink modules are age-dependent modules that show similar changes in protein abundance in control and rNLS8 mice, which may be attributed to the age of the mice at the timepoints analyzed. The number of proteins in each module is listed in parentheses. *Left,* Log fold change of module proteins in control and rNLS8 samples at each disease stage. *Center*, Heat map of module proteins. *Right,* Gene ontology of the biological processes enriched in each module. The size of the circles is dependent on enrichment ratio and the top 5 proteins belonging to the most significant term are shown ranked by kMe value (module membership score) for **b,** green (ribosome), **c,** red (mitochondria), and **d,** pink (cytoskeletal organization) modules.

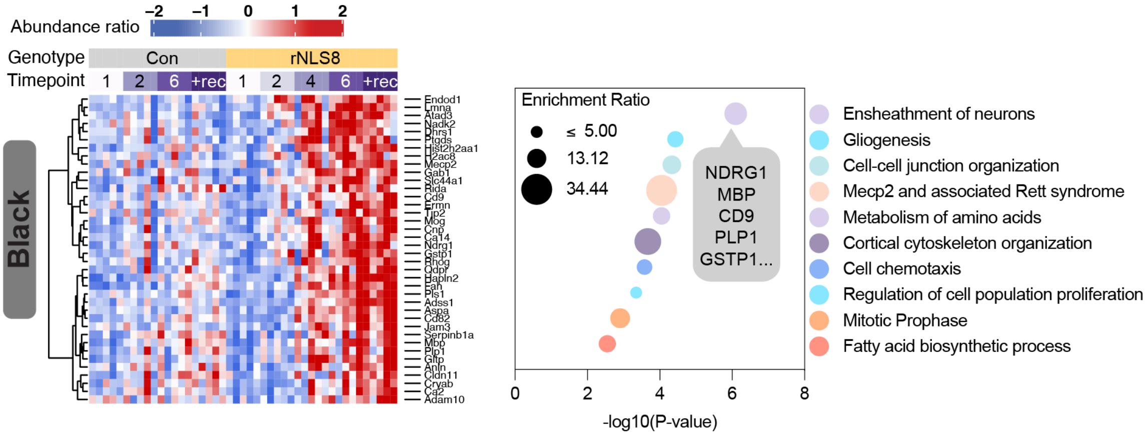


**Supplementary Figure 2. Gene ontology analysis of the black module identifies myelin ensheathment of neurons and gliogenesis as key increased pathways in the rNLS8 cortex.** Proteins in the black module showed a time-dependent increase in abundance in rNLS8 mouse cortex but were stable across time in control animals (Figure 3c). The abundance of these proteins remained high in rNLS8 cortex in recovery, suggesting that disease-associated changes in this module may not directly correlate with TDP-43 pathology burden, and these alterations may be required for initiation of motor recovery after inhibition of hTDP-43ΔNLS expression. Proteins in the black module were enriched in proteins expressed highly in oligodendrocytes, the myelinating glia of the central nervous system (enrichment ratio = 4.79, $P < 0.00001$; Figure 3g). Complex heatmap of protein abundance ratios (*left*) of each protein identified in the black module and the top 10 gene ontology terms plotted by significance (*right*). Overlaid with callouts that list 5 proteins that comprise the top biological process in each module ranked by kME value.

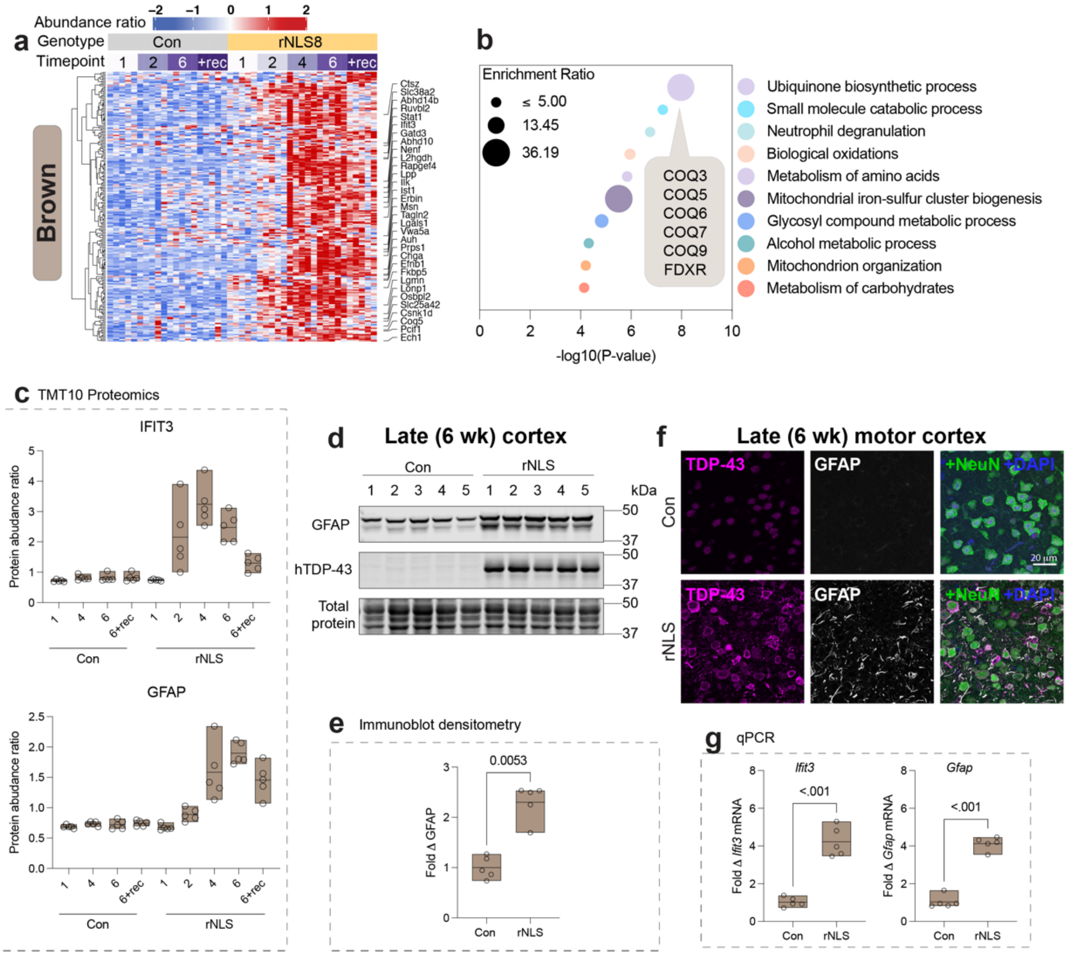


**Supplementary Figure 3. Gene ontology analysis revealed that the brown module was enriched in ubiquinone biosynthesis and small molecule catabolic processes that are increased in late disease and are partially reversed with recovery in rNLS8 mice.** Proteins in the brown module showed a time-dependent increase in abundance in rNLS8 mice beginning in later disease stages, with no change over time in control mice. The elevated abundance of these proteins was partially reversed in rNLS8 recovery mice compared to controls. The brown module was enriched for microglial proteins (enrichment ratio = 2.05, $P < 0.05$) (Figure 3g). **a,** Heatmap of protein abundance of all proteins in the brown (catabolism) module. **b,** Gene ontology of enriched biological processes in the brown module. Ranked by P-value and circle size is dependent on enrichment ratio. **c,** Protein abundance ratio from quantitative temporal proteomics of interferon-induced protein with tetratricopeptide repeats 3 (IFIT3) and glial fibrillary acid protein (GFAP). IFIT3, an antiviral protein expressed in response to interferon-alpha, and GFAP, a marker of astrogliosis, exhibited the highest fold changes in the brown module. **d,** Immunoblot of RIPA-soluble whole cortex tissue lysates after 6 weeks of cytoplasmic TDP-43 expression. Tissue was probed for GFAP (50 kDa) and human TDP-43 (43 kDa). **e,** Immunoblot densitometry, fold change protein levels relative to the mean of control mice and normalized to total protein loading. **f,** Immunofluorescence microscopy of the primary motor cortex in control and rNLS8 mice after 6 weeks of cytoplasmic TDP-43 expression. Samples were immunolabeled for GFAP (*top to bottom*), and co-labeled for TDP-43, NeuN (pan-neuronal marker) and DAPI. Scale bar = 20 μm. **g,** Fold change in mRNA transcript level of target genes relative to the mean of control mice and normalized to GAPDH housekeeping gene by qPCR. Antibodies tested against IFIT3 were ineffective; however, we detected a 4.2-fold increase in *Ifit3* mRNA in cortex tissue from mice in late disease, suggesting that the brown module represents transcriptionally mediated inflammatory activation. GFAP demonstrated an increase in mRNA (4.1-fold increase), protein levels by immunoblot (2.3-fold increase), and astrocytic localization by immunohistochemistry in late-disease cortex tissue. All data is from n = 5 mice per group (open circle points) with a line at the mean and range displayed by floating bars. Differences in the means of control and rNLS8 mice were determined using a two-tailed paired t-test, where a value of $P < 0.05$ was considered statistically significant.

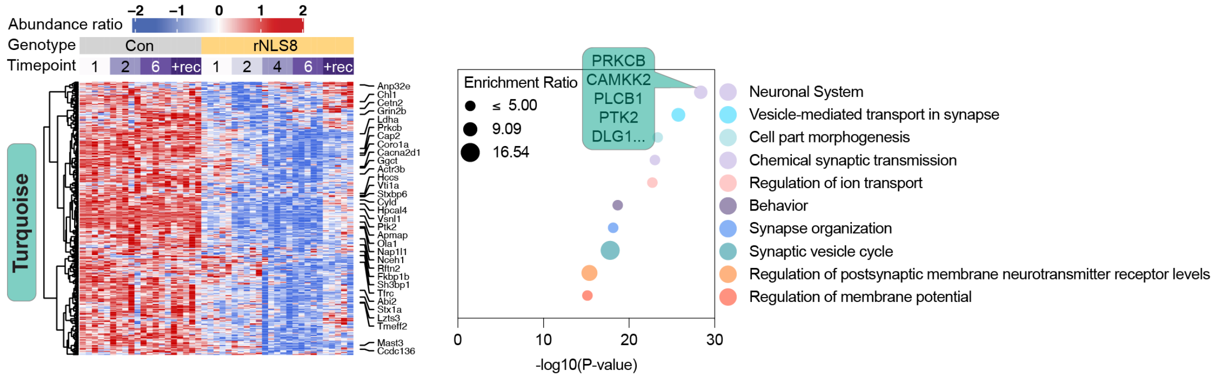


**Supplementary Figure 4. Gene ontology of protein subsets that comprise the turquoise module identifies the neuronal system as a key decreased pathway in disease.** Complex heatmaps of protein abundance ratios (*left*) of each protein identified in the turquoise module and the top 10 gene ontology terms plotted by significance (*right*). Overlaid with callouts that list 5 proteins that comprise the top biological process in each module ranked by kME value.

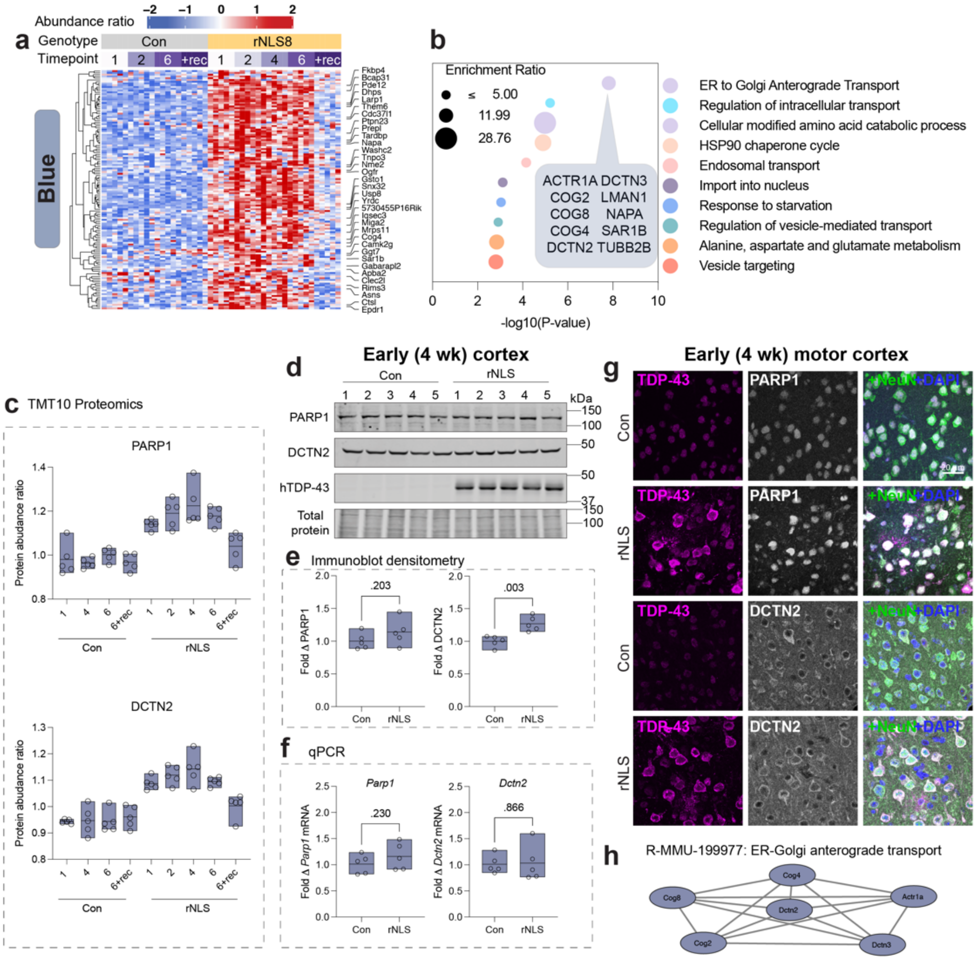


**Supplementary Figure 5. Proteins involved in transport in the blue module demonstrate early and sustained altered levels in the rNLS8 cortex that return to control levels in recovery.** The blue module was characterized by early increases in abundance at pre-onset (1 wk) that were sustained throughout the disease course in the rNLS8 cortex, with no change over time in control mice (Figure 3c). The abundance of proteins in the blue module returned to control levels in recovery and correlated with levels of hTDP-43ΔNLS. **a,** Heatmap of protein abundance of all proteins in the blue module. **b,** Gene ontology of enriched biological processes in the blue module revealed that ER to Golgi anterograde transport and regulation of intracellular transport were the top two significantly enriched biological processes. Ranked by P-value, with circle size being dependent on enrichment ratio. **c,** Protein abundance ratio from quantitative temporal proteomics of Poly [ADP-ribose] polymerase 1 (PARP1) and dynactin subunit 2 (DCTN2). **d,** Immunoblot of RIPA-soluble whole cortex tissue lysates after 4 weeks of cytoplasmic TDP-43 expression. Tissue was probed for PARP1 (115 kDa), DCTN2 (50 kDa), and human TDP-43 (43 kDa). **e,** Immunoblot densitometry, fold change protein levels relative to the mean of control mice and normalized to total protein loading. **f**, Fold change in mRNA transcript level of target genes relative to the mean of control mice and normalized to the GAPDH housekeeping gene by qPCR. Immunoblotting and immunohistochemistry validated the increase in DCTN2 in rNLS8 mice compared to controls; however, there was no change in *Dctn2* transcript level, indicating that protein level increases were not due to increased gene expression. All data are from n = 5 mice per group (open circle points) with a line at the mean and range displayed by floating bars. Differences in the means of control and rNLS8 mice were determined using a two-tailed paired t-test, where a value of $P < 0.05$ was considered statistically significant. **g,** Immunofluorescence microscopy of the primary motor cortex in control and rNLS8 mice in early disease (4 wk). Samples were immunolabeled for each target, PARP1 and DCTN2 (*top to bottom*), and co-labeled for TDP-43, NeuN (pan-neuronal marker) and DAPI. Scale bar = 20 μm. Immunohistochemistry validated the increase in PARP1 in the nucleus of neurons in the primary motor cortex of rNLS8 mice compared to controls, but there was no significant change in *Parp1* at the transcript level or by immunoblot. **h,** Protein-protein interaction network for proteins, including DCTN2, in the blue module that are involved in ER to Golgi anterograde transport.

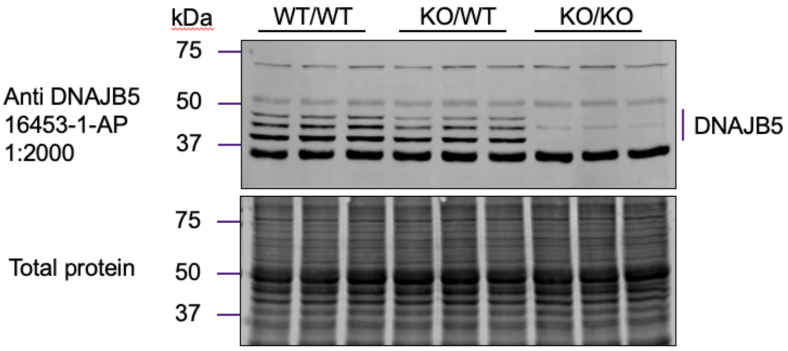


**Supplementary Figure 6. Validation of DNAJB5 antibody in *Dnajb5*$^{KO/KO}$ knockout mice.** RIPA-soluble cortex tissue from wild-type (WT/WT), heterozygous DNAJB5 knockout (KO/WT) and homozygous DNAJB5 knockout (KO/KO) mice (n = 3) were probed for DNAJB5 using the Proteintech 16453-1-AP antibody. The antibody detected multiple bands between 37-75 kDa, three of which were eliminated or decreased with DNAJB5 KO that represent DNAJB5 isoforms in the cortex.

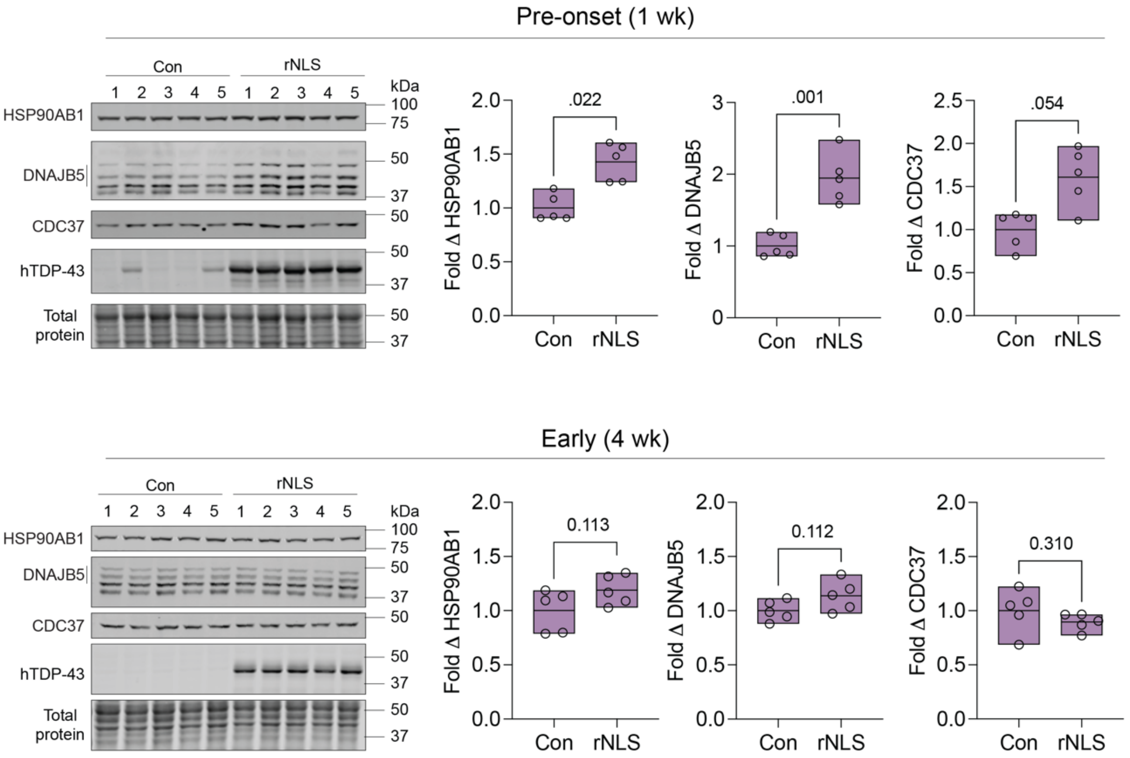


**Supplementary Figure 7. Magenta module proteins, HSP90AB1, DNAJB5 and CDC37 are increased at pre-onset and disease onset but normalize to control levels in early disease.** Immunoblot analysis of control and rNLS8 mouse cortex RIPA-soluble lysates at pre-onset (1 wk; top) and in early disease (4 wk; bottom). Samples were probed for HSP90AB1, DNAJB5, CDC37, TDP-43, and Revert total protein stain and the relative abundance of each protein was normalized to the total protein loading. Data are presented as open circles to represent individual mice (n = 5/group), and floating bars show the mean and range of the data. Differences between the means were determined using a paired t-test, where P<0.05 was considered statistically significant.

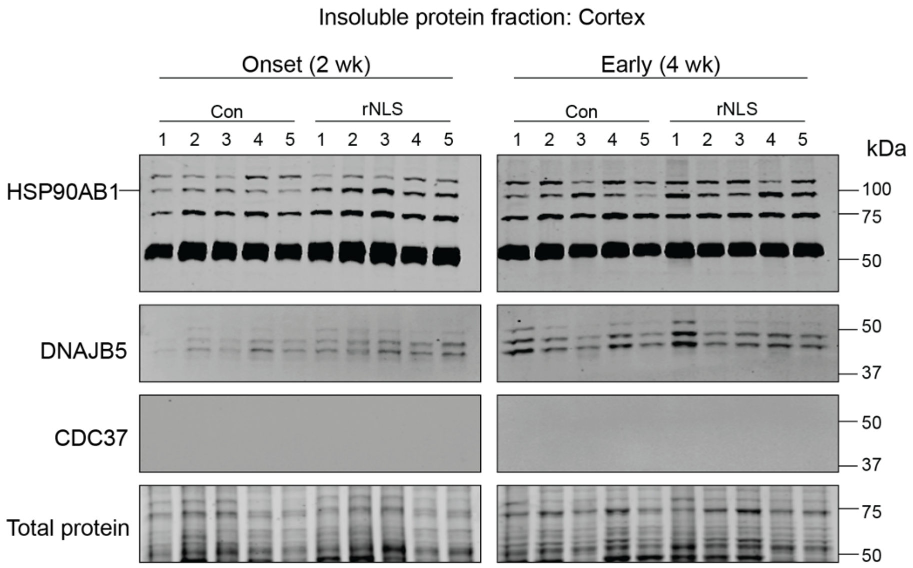


**Supplementary Figure 8. Not all chaperone proteins in the magenta module are sequestered to the insoluble protein fraction in the cortex of rNLS8 mice.** The cortex of rNLS8 mice at onset (2 wk) and in early disease (4 wk) was harvested and fractionated into soluble and insoluble fractions. The insoluble protein fraction was immunoblotted and probed for HSP90AB1, DNAJB5, and CDC37 to determine whether there was sequestration of these proteins into the insoluble fraction. We observed HSP90AB1 and DNAJB5 in the insoluble fraction of control and rNLS8 cortex tissue at disease onset and early disease, but there was no insoluble CDC37 at either timepoint.

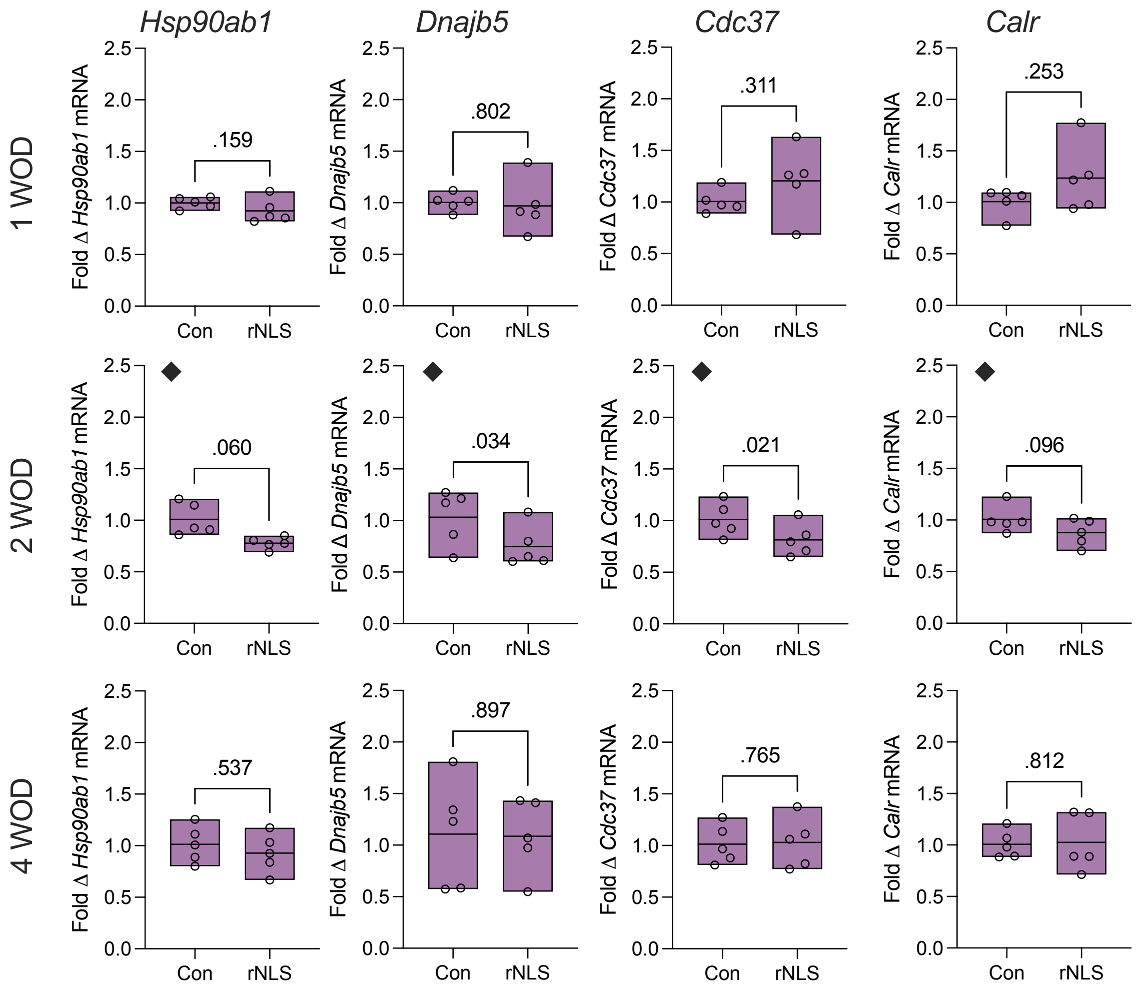


**Supplementary Figure 9. Transcript levels of magenta module proteins are not significantly increased in the cortex of rNLS8 mice, despite increased protein levels at pre-onset and onset shown in Figure 4 and Supplementary Figure 8.** ◆ Denotes data that was shown in Figure 4, which are shown again here for a complete dataset of the transcript levels at each timepoint, pre-onset (1 wk), onset (2 wk) and early disease (4 wk). Data are shown as the fold change in mRNA transcript level of target genes relative to the mean of control mice and normalized to the GAPDH housekeeping gene by qPCR. All data are from n = 5 mice per group (open circle points) with a line at the mean and range displayed by floating bars. Differences in the means of control and rNLS8 mice were determined using a two-tailed paired t-test, where $P < 0.05$ was considered statistically significant.

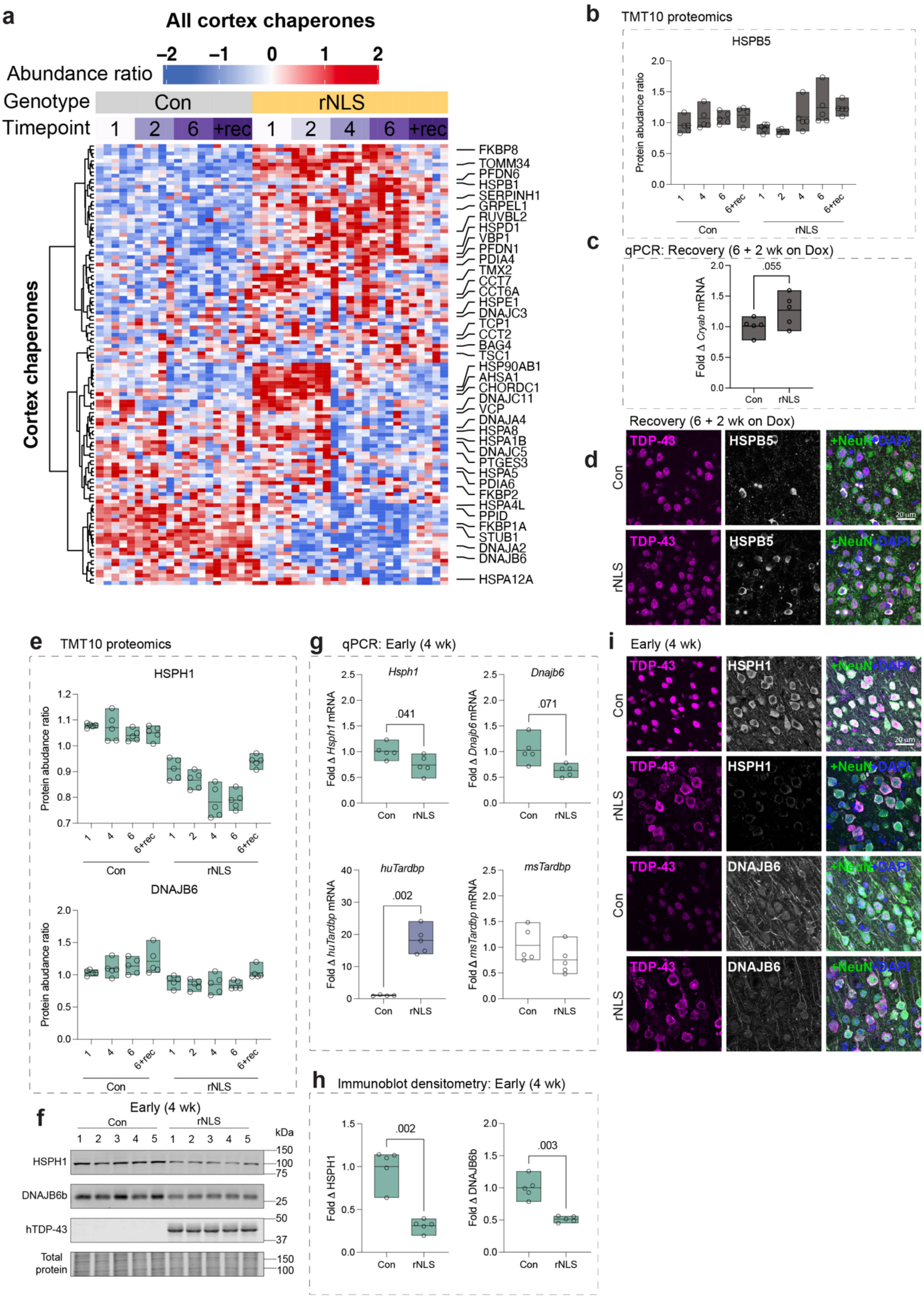


**Supplementary Figure 10. Chaperone proteins show diverging patterns of abundance in the rNLS8 cortex. a,** Heatmap of a subset of the longitudinal proteomics data representing 119 chaperones identified in the cortex. Samples are from control and rNLS8 mice 1, 2, 4, 6 weeks off dox and recovery mice (6 weeks off dox + 2 weeks on dox). Red is high and blue is low relative protein abundance. **b-d,** Analysis of cortex tissue from recovery mice (6 wk off dox + 2 weeks on dox) **b,** Relative protein abundance of HSPB5 from quantitative longitudinal proteomics. **c,** Fold change in mRNA transcript level of *Cryab* (which codes for HSPB5) relative to the mean of control mice and normalized to the *Gapdh* housekeeping gene by qPCR. **d,** Immunofluorescence microscopy of the primary motor cortex in control and rNLS8 mice in recovery shows clearance of cytoplasmic TDP-43 and nuclear localization of TDP-43. Samples were immunolabeled for HSPB5, and co-labeled for TDP-43, NeuN (pan-neuronal marker), and DAPI. **e,** Protein abundance ratio from quantitative temporal proteomics of heat shock protein family H member 1 (HSPH1), and DnaJ heat shock protein family member (Hsp40) member B6 (DNAJB6). **f,** Immunoblot of RIPA-soluble whole cortex tissue lysates after 4 weeks of cytoplasmic TDP-43 expression. Tissue was probed for HSPH1 (105 kDa), DNAJB6b (26 kDa), and human TDP-43 (43 kDa). **h,** Immunoblot densitometry, fold change protein levels relative to the mean of control mice and normalized to total protein loading. **g,** Fold change in mRNA transcript level of target genes relative to the mean of control mice and normalized to the *Gapdh* housekeeping gene by qPCR. **i,** Immunofluorescence microscopy of the primary motor cortex in control and rNLS8 mice in early disease. Samples were immunolabeled for each target, HSPH1 and DNAJB6 (*top to bottom*), and co-labeled for TDP-43, NeuN (pan-neuronal marker), and DAPI. All data shown are from n = 5 mice per group (open circle points) with a line at the mean and range displayed by floating bars. Differences in the means of control and rNLS8 mice were determined using a two-tailed paired t-test, where a value of $P < 0.05$ was considered statistically significant. Scale bar = 20 μm.

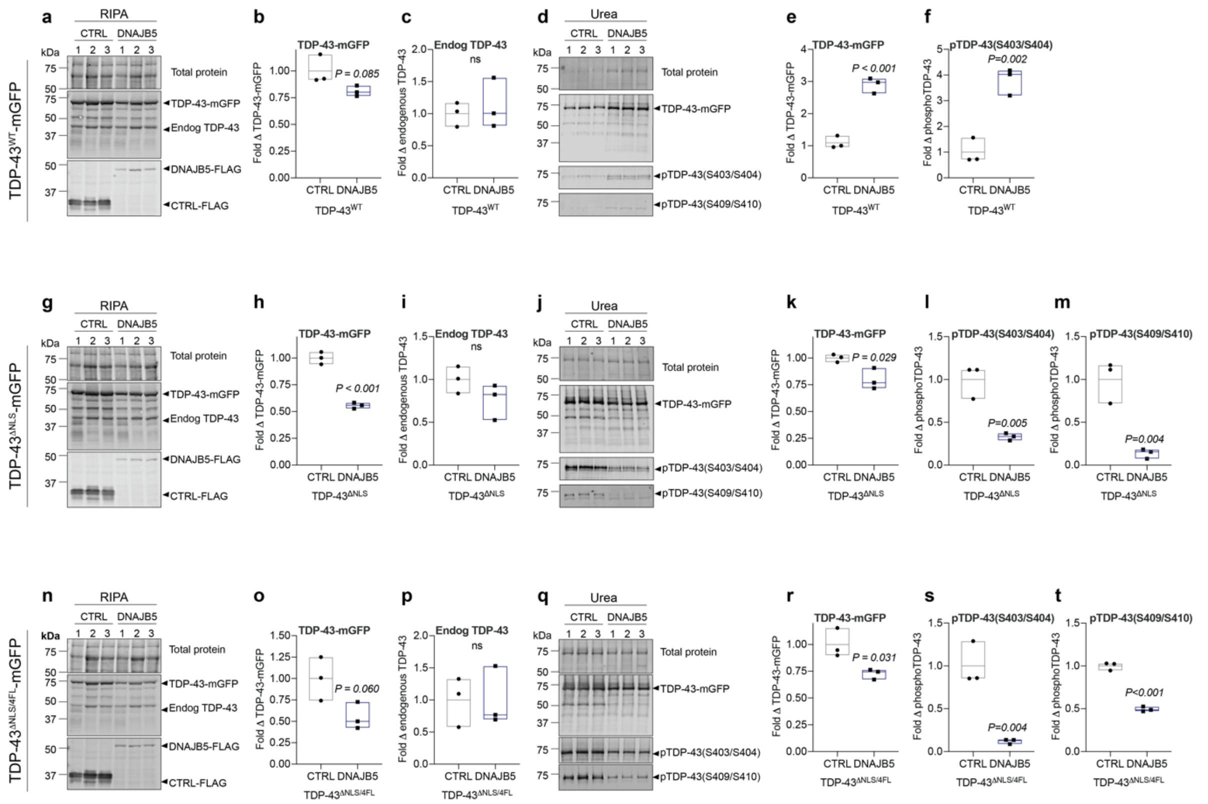

**Supplementary Figure 11. Over-expression of DNAJB5 decreases soluble and insoluble mutant TDP-43 in HEK293 cells.** HEK293 cells were co-transfected to express wild-type or mutant forms of TDP-43-mGFP and DNAJB5-FLAG or a negative control (CTRL-FLAG). Cell lysates were fractionated into RIPA-soluble and urea-soluble fractions for immunoblotting. **a**, Immunoblot of RIPA-soluble TDP-$43^{WT}$-mGFP and densitometry of **b**, TDP-43, and **c**, endogenous TDP-43. **d**, Immunoblot of urea-soluble TDP-$43^{WT}$ and densitometry of **e**, TDP-$43^{WT}$-mGFP and **f**, phospho-TDP-43 (S403/S404). **g**, Immunoblot of RIPA-soluble TDP-$43^{\Delta NLS}$-mGFP and densitometry of **h**, TDP-$43^{\Delta NLS}$-mGFP, and **i**, endogenous TDP-43. **j**, Immunoblot of urea-soluble TDP-43 and densitometry of **k**, TDP-$43^{\Delta NLS}$-mGFP, **l**, phospho-TDP-43 (S403/S404), and **m**, phospho-TDP-43 (S409/S410). **n**, Immunoblot of RIPA-soluble TDP-43 and densitometry of **o**, TDP-$43^{\Delta NLS/4FL}$-mGFP, and **p**, endogenous TDP-43. **q**, Immunoblot of urea-soluble TDP-43 and densitometry of **r**, TDP-$43^{\Delta NLS/4FL}$-mGFP, **s**, phospho-TDP-43 (S403/S404), and **t**, phospho-TDP-43 (S409/S410). Statistically significant differences between the means were determined using unpaired t-tests, where $P < 0.05$ was considered significant.

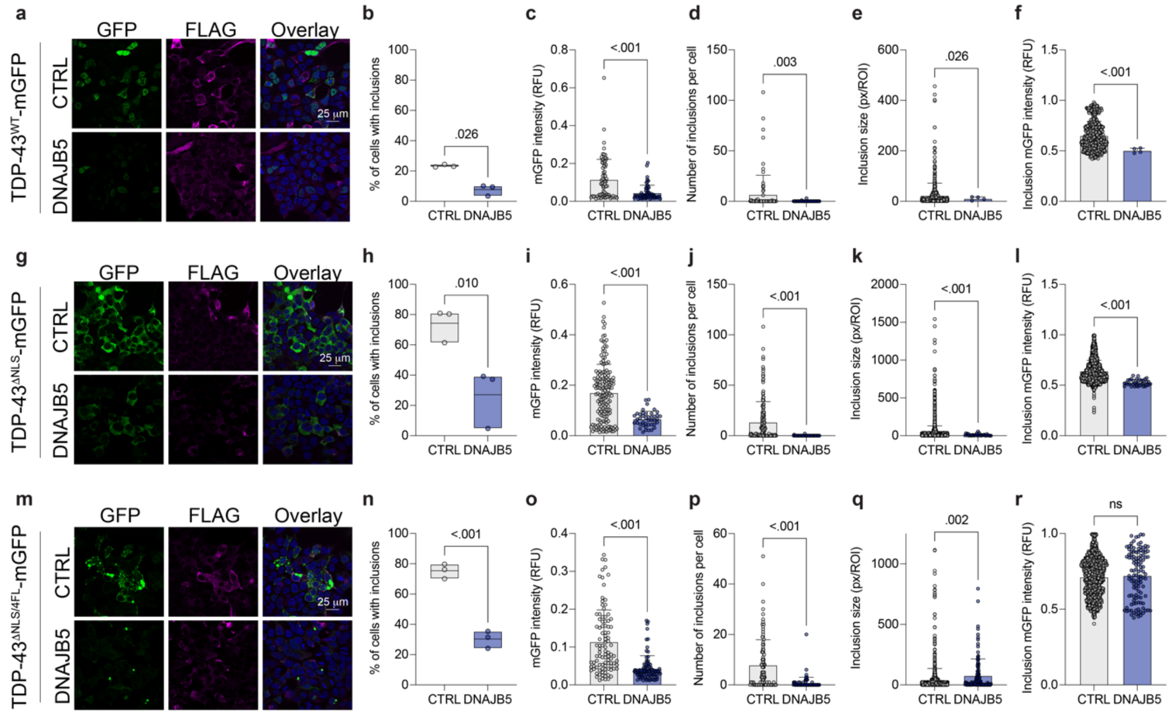


**Supplementary Figure 12. Over-expression of DNAJB5 decreases the proportion of cells with mutant TDP-43 inclusions and the number of inclusions per cell.** HEK293 cells were co-transfected to express wild-type or mutant forms of TDP-43-mGFP and DNAJB5-FLAG or a negative control (CTRL-FLAG). Cells were fixed and immunolabeled with anti-FLAG, imaged by microscopy, and subjected to non-biased single cell analysis using Cell Profiler. Cells over-expressing **a**, TDP-43$^{WT}$-mGFP, **g**, TDP-43$^{\Delta NLS}$-mGFP, or **m**, TDP-43$^{\Delta NLS/4FL}$-mGFP and CTRL or DNAJB5 were analyzed to determine **b/h/n**, the percent of cells with TDP-43WT-mGFP inclusions, **c/i/o**, fluorescence intensity of mGFP in relative fluorescence units (RFU), **d/j/p**, number of inclusions per cell, **e/k/q**, inclusion size, and **f/l/r**, inclusion mGFP fluorescence intensity. Statistically significant differences between the means were determined using unpaired Welch's t-tests, where P < 0.05 was considered significant.

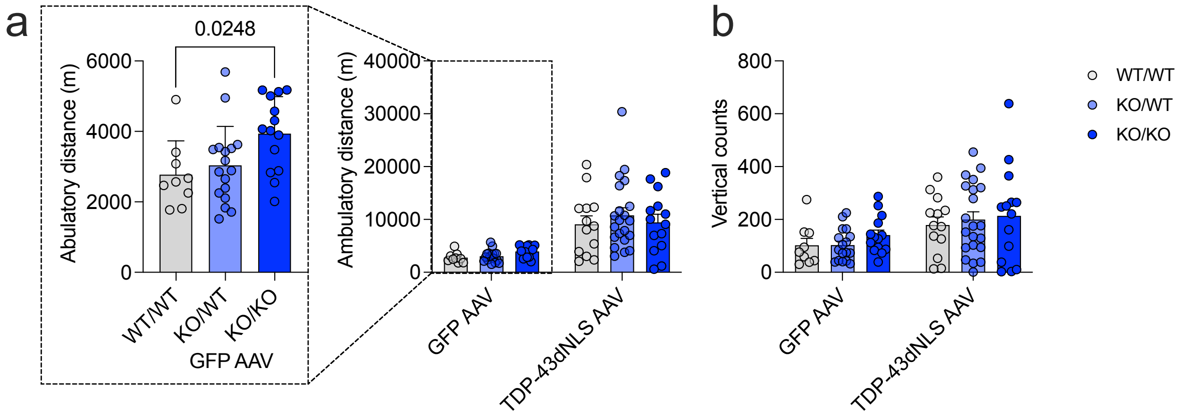


**Supplementary Figure 13. Open field behaviour testing of *Dnajb5* knockout mice injected with TDP-43DNLS-myc or GFP-myc (control) AAVs.** *Dnajb5* wildtype (WT/WT), heterozygous (KO/WT), and homozygous *Dnajb5* knockout (KO/KO) P0 pups were injected with human cytoplasmic TDP-43DNLS-myc or GFP-myc (control) AAVs via bilateral intracerebroventricular injections. Mice at 12 weeks of age were subjected to open field testing to assess their relative **a**, ambulatory distance (in meters) and **b,** vertical counts. Data shown is the mean + standard error of the mean and data points represent individual mice (*n* = 10-21/group).

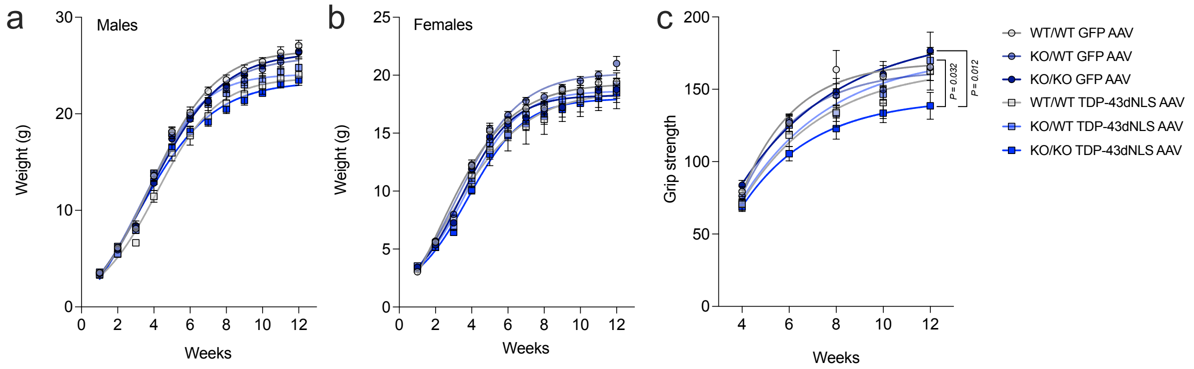


**Supplementary Figure 14. Weight and grip strength testing of *Dnajb5* knockout mice injected with TDP-43DNLS-myc or GFP-myc (control) AAVs.** *Dnajb5* wildtype (WT/WT), heterozygous (KO/WT), and homozygous *Dnajb5* knockout (KO/KO) P0 pups were injected with human cytoplasmic TDP-43DNLS-myc or GFP-myc (control) AAVs via bilateral intracerebroventricular injections. Mice were subjected to weekly weighing and grip strength testing every other week. **a**, Weight in grams of males over time. **b,** Weight in grams of females over time. **c**, Grip strength (gram force) of males and females combined over time. Data shown is the mean + standard error of the mean and data points represent individual mice (*n* = 10-21/group).

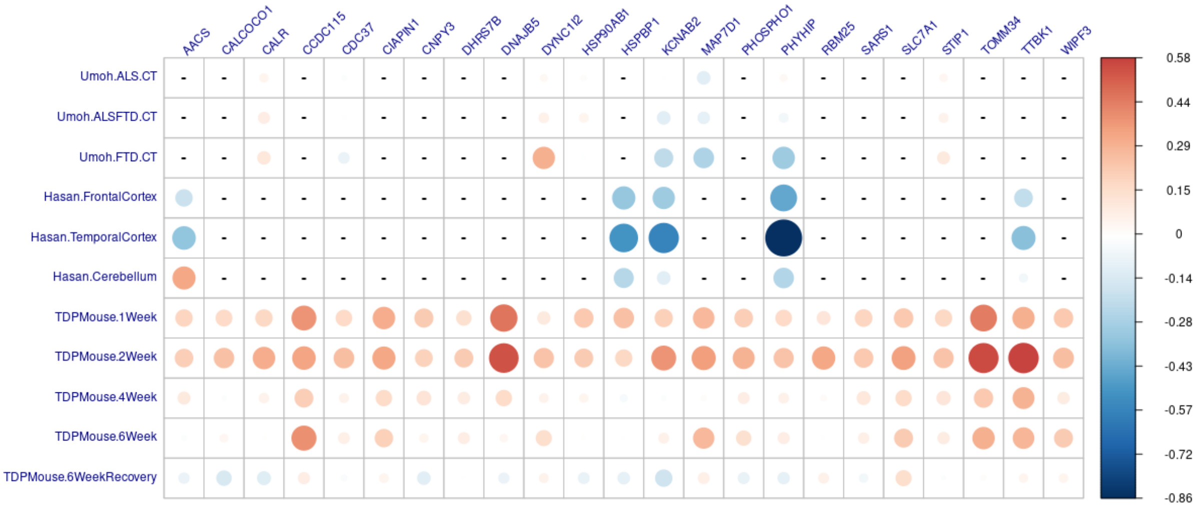


**Supplementary Figure 15. Protein/transcript abundance (log fold change) of protein folding magenta module proteins in human postmortem TDP-43 proteinopathies and the rNLS8 mouse cortex.** Proteins/transcripts for magenta module proteins are listed on the top and define each column. Each comparison made within each study is listed on the y-axis where each row of the table represents the log fold change of each protein/transcript in each comparison. Circle size is dependent on the magnitude of change and circle colour is dependent on log fold change where blue is decreased and red is increased compared to controls.
